# The development and characterization of an scFv-Fc fusion based gene therapy to reduce the psychostimulant effects of methamphetamine abuse

**DOI:** 10.1101/687129

**Authors:** Charles E. Hay, Laura E. Ewing, Michael D. Hambuchen, Shannon M. Zintner, Juliana C. Small, Chris T. Bolden, William E Fantegrossi, Paris Margaritis, S. Michael Owens, Eric C. Peterson

## Abstract

Methamphetamine (METH) continues to be amongst the most addictive and abused drugs in the US. Unfortunately, there are currently no FDA approved pharmacological treatments for METH substance abuse disorder. As an alternative approach, we have previously explored the use of Adeno-associated viral (AAV) mediated gene transfer of an anti-METH monoclonal antibody. Here, we advance our approach by generating a novel anti-METH scFv-Fc fusion construct (7F9-Fc), packaged into AAV serotype 8 vector (called AAV-scFv-Fc), and tested in vivo and ex vivo. A range of doses (1 × 10^10^. 1 × 10^11^, and 1 × 10^12^ vector copies(vc)/mouse) were administered to mice, which exhibited a dose-dependent expression of 7F9-Fc in serum with peak circulating concentrations of 48, 1785, and 3,831 μg/ml. The dose of 1 × 10^12^ vc/mouse was further tested in METH locomotor and biodistribution studies to determine the efficacy of the antibody protection. Expressed 7F9-Fc exhibited high affinity binding, 17 nM, to METH. Between days 21 and 35 after vector administration, the 7F9-Fc gene therapy significantly reduced the effects of METH in locomotor assays following administration of moderate and high doses of subcutaneous METH, 3.1 and 9.4 mg/kg respectively. On day 116 post-AAV administration, mice expressing 7F9-Fc sequestered over 2.5 times more METH into the serum than vehicle mice, and METH concentrations in the brain were reduced by 1.2 times compared to vehicle mice. Taken together, these data suggest that a AAV-delivered anti-METH Fc fusion antibody could be a design for persistently reducing concentrations of METH in the CNS.

**Significance Statement:** In this manuscript, we describe the use of a novel anti-METH scFv-Fc fusion protein delivered in mice using gene therapy. The results suggest that the gene therapy delivery system can lead to the production of enough antibody to mitigate METH’s psychostimulant effects in mice over an extended time period.

## Introduction

METH abuse has been increasing yearly since at least 2008 and its use as an illicit drug is second only to opioid abuse (Gorman, 2018; Artigiani *et al.*, 2019). The United States has seen a 470% increase in hospitalizations involving METH abuse in the last seven years (Winkelman *et al.*, 2018). While the accepted treatment for METH addiction recovery consists primarily of 8-12 months of behavioral therapy sessions (Grant *et al.*, 2012), the lack of an FDA approved pharmacological therapy hinders recovery. Without a pharmacological component of treatment, recovery from METH addiction is difficult, with rates of recidivism over 60% within the first year of abstinence (Brecht and Herbeck, 2014).

One pharmacological approach is to use anti-METH monoclonal antibodies (mAbs) as pharmacokinetic antagonists to block or blunt METH effects. Unlike a small molecule medication, which would selectively block multiple receptors for extended periods of time, mAbs sequester METH in the circulatory system (Stevens, Henry, *et al.*, 2014; Stevens, Tawney, *et al.*, 2014; Hay *et al.*, 2018) and reduce METH entry into critical tissues like the central nervous system (CNS) and the cardiovascular system. Pharmacokinetic assays demonstrated that an anti-METH mAb therapy, administered as a bolus *i.v.* injection, has an apparent half-life of about three weeks in humans and one week in rats (Stevens, Henry, *et al.*, 2014; Stevens, Tawney, *et al.*, 2014).

Delivery of the mAb by gene therapy could greatly extend the duration of action, reduce dosing frequencies, and avoid potential noncompliance issues. Adeno-associated viruses (AAV) is commonly used as gene therapy vectors, because it is a non-pathogenic virus that can deliver up to 4.5 kilobases of DNA to host tissues (Zolotukhin, 2005). Previous studies have shown that a single dose of AAV, encoding an entire mAb sequence, reaches maximal expression of mAb in 3-4 weeks and expression persists for at least 8 months (Hicks *et al.*, 2012; Rosenberg *et al.*, 2012, 2013; Hay *et al.*, 2018). Hicks *et al.* (2012) showed that an anti-nicotine AAV based mAb therapy elicited over 4-fold greater circulating concentrations of high affinity mAbs with one injection compared to an anti-nicotine vaccine, which generated variable amounts of polyclonal antibodies and required one to two additional booster doses. These previous studies suggest that an optimized anti-METH gene therapy should be able to sequester greater levels of METH in the serum than a METH vaccine could.

Other labs have successfully developed AAV-based mAb therapies against various drugs of abuse (Hicks *et al.*, 2012; Rosenberg *et al.*, 2012). A full IgG antibody sequence can fit into an AAV capsid, but requires post-translational autocleavage of the heavy and light chains of the IgG antibody before dimerizing back together through disulfide bonds (Fang *et al.*, 2005; Ho *et al.*, 2013). In contrast, we have previously reported the design and testing of a singlechain variable fragment (scFv), which consists of only the variable regions of an IgG connected by a 50 amino acid linker (Hay *et al.*, 2018). Because scFvs have a short half-life of roughly 60 min, limited circulating concentrations of anti-METH antibodies were observed in mice (Nanaware-Kharade *et al.*, 2015; Reichard *et al.*, 2016; Hay *et al.*, 2018).

A compromise was discovered in an scFv-Fc fusion protein, which consists of an scFv, connected to the C_H2_ and C_H3_ regions of an IgG constant region (Unverdorben *et al.*, 2016). This protein can then create a homodimer with another copy of itself. An scFv-Fc fusion has the potential to avoid the more complex post-translational processing that is necessary for IgG assembly, while still maintaining a long half-life. ScFv-Fc fusions are reported to have serum half-lives of 104-112 hrs compared to the 218-222 hrs of a full IgG antibody (Unverdorben *et al.*, 2016).

This leads to the hypothesis that the inclusion of the Fc region from an IgG antibody, forming an scFv-Fc fusion antibody against METH, would yield high circulating concentrations. These high concentrations of the novel antibody should sequester greater concentrations of METH in the circulation and significantly reduce the psychostimulant effects of METH in mice.

In this study, we report a newly designed anti-METH scFv-Fc biotherapeutic (7F9-Fc) packaged into an AAV8 viral vector. We report AAV8 viral vector dose-dependent 7F9-Fc serum concentrations in mice that show antibody circulating concentrations in the mg/ml range over a 7 month period. Studies of locomotor activity showed that METH psychostimulant effects were reduced at drug doses up to 10 mg/kg, and biodistribution studies showed that there were significant increases in the METH brain:serum ratio at 30 min after a 3.1 mg/kg METH dose.

## Methods

### Animal usage

Adult (3-4 weeks old), male, BALB/c mice were ordered from Charles River Laboratories (Raleigh, NC). Mice were housed with 3-6 mice per cage in a light-controlled environment (12 hr light/dark cycle). Mice were allowed food and water *ad libitum.* Mice were randomly placed into groups for all experimental studies. The use of mice for experiments was approved by the Institutional Care and Use Committee of the Universities of Arkansas for Medical Sciences and was in accordance with the “Guide for the Care and Use of Laboratory Animals” (National Research Council 2011).

### Drugs and Reagents

All (+)-methamphetamine (METH) used, both unlabeled and tritium-labeled, was obtained from the National Institute of Drug Abuse (Bethesda, MD). Unless otherwise stated, reagents and other materials were purchased from ThermoFisher Scientific (Waltham, MA). Plasmid and protein isolation and purification was performed using Qiagen kits (Valencia, CA). METH ampules were bought for standard curve samples and quality control samples from Cerilliant (Round Rock, TX). DNA digestion enzymes were purchased from New England Biolabs (Ipswich, MA).

### Design and generation of scFv-Fc plasmids and AAV8 capsids

The scFv-Fc fusion construct, 7F9-Fc, was derived from an anti-METH mAb previously generated, consisting of scFv7F9 and C_H2_ and C_H3_ regions (Peterson *et al.*, 2008). The sequence was capped with a 6-histidine tail (6HIS) at the 3’ end, and expressed as a cleavable, signal secretion sequence (HMM38) to ensure correct folding and routing through the cellular secretory pathway (Barash *et al.*, 2002). Plasmids were cloned into cassettes with a a human a-1 anti-trypsin promoter (HAAT) prior to the HMM38 sequence. The cassette structure is shown in Figure 1A.

**Figure 1.**
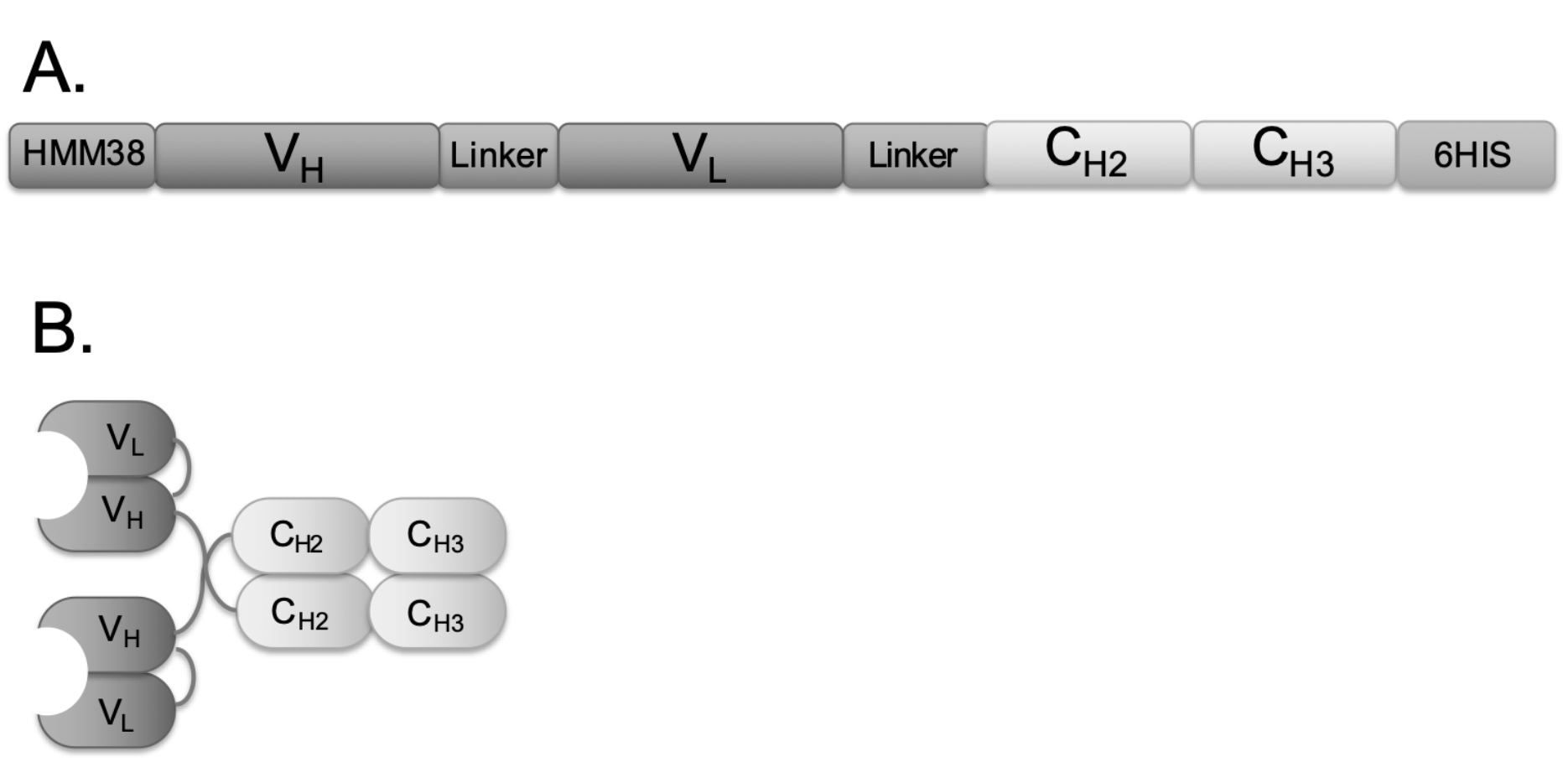
Schematic of the scFv-Fc fusion construct. 1A shows a schematic of the different regions of the DNA sequence packaged into the AAV capsids. 1B shows the same sequences after being translated and folded; construct should form a homodimer before being secreted from the cell. HMM38, secretory signal; V_H_, variable heavy region; Linker, amino acid linker; V_L_, variable light region; C_H2_ and C_H3_, constant regions 2 and 3 respectively; and 6His, 6-histidine tag for purification and identification.

The sequences containing 7F9-Fc cDNA were custom synthesized and ligated into pUC57 (GenScript). Plasmids containing 7F9-Fc were transformed into Invitrogen Top10F competent cells (Thermo #C303003) as per manufacturer recommendations and sequences were confirmed by sequencing at the University of Arkansas for Medical Sciences DNA Sequencing Core Facility. Plasmid were transformed into Top10F cells and colonies of Top10F cells were grown on agar plates containing 50 μg/ml ampicillin (AMP). Colonies were selected and added to baffled flasks containing 1 liter of LB with 50 μg/ml ampicillin (AMP). Cultures were grown up overnight with shaking at 225 rpm at 37° C. Qiagen Endotoxin-free Mega (#12381) plasmid kits were used to isolate the plasmid DNA. DNA purity and concentrations were determined by UV_260/280_ spectrophotometry and gel electrophoresis via plasmid digestions using the digestion enzymes EcoRI and XhoI (New England Biolabs # R3101S and R0146S respectively).

Plasmids containing the construct DNA were shipped to the Children’s Hospital of Philadelphia to be ligated into the expression vector, AAV–hAAT-F.IX as described in Margaritis *et al.*, (2004). These ligated expression vectors were then sent to SAB Tech (Philadelphia, Pennsylvania), who employs a helper virus-free, triple plasmid transfection method for AAV8 vector production and packaging into viral vectors.

### Determining Titers and Duration of Expression

Four to six mice per group were administered 1 × 10^10^, 1 × 10^11^, or 1 × 10^12^ vector copies (vc) of AAV-7F9-Fc per mouse or equal volumes of 5% sorbitol (w/v) in 1X PBS for the vehicle group via tail vein. Tail snips were performed every two to four weeks to collect blood samples from each mouse (Hay *et al.*, 2018). The serum was isolated from the red blood cells via centrifugation (12,000 rcf for 10 min) and stored at −80° C.

Functional, sandwich ELISAs were performed as previously described to determine the circulating concentrations of 7F9-Fc at each time point (Hay *et al.*, 2018). Briefly, the serum samples were diluted 1:5,000, and a HRP anti-His tag antibody (Biolegend, #652504) diluted 1:1000 in Superblock (Thermo #37515) was used as the secondary detection antibody. 1-Step Ultra TMB-ELISA (Thermo #34029) was utilized to react with the HRP on the secondary antibodies. Wells were measured for light absorption at 480 nm on a Biotek HT Synergy plate reader. Concentrations of the anti-METH antibodies were interpolated from a standard curve of purified scFv7F9 protein (Reichard *et al.*, 2016).

### Biocharacterization

Western blots were performed as previously described to confirm the relative size of the expressed 7F9-Fc constructs (Hay *et al.*, 2018). Briefly, serum samples collected 22 days post-AAV administration were pooled from each AAV dose group and diluted 1:13 before applying to the Western blot membranes. Serum pooled from vehicle mice on day 22 was used as a negative control to show mice not given the AAV-7F9-Fc vectors did not express 7F9-Fc protein. Purified scFv7F9 protein with a 6His tag was used as a positive control (Reichard *et al.*, 2016). The Western blot membranes were stained with HRP anti-His tag antibody (Biolegend, #652504) diluted 1:2,000 in 0.5% (w/v) BSA. Membranes were developed using Pierce ECL Western Blotting Substrate (Thermo #32106). Due to the use of a standard curve in the ELISA assays to precisely interpolate expression level, western blot bands were not quantified.

### Determination of AAV-7F9-Fc antibody ex vivo function

Radioimmunoassays were performed as previously described to determine the IC_50_ values for METH binding to 7F9-Fc (Owens *et al.*, 1988; Peterson *et al.*, 2007). Briefly, 10 μl of serum samples or METH serum standands (concentrations from 0.03 nM to 500 nM) were added to duplicate polypropylene tubes containing: 20 μl of magnetic MagnaBind protein G beads (Thermo #213498), 100 μl of 500 dpm/μl ^3^H-METH, and 100 μl of blank serum diluted 1:1000. After 18 hrs of gentle mixing at 4° C, the tubes were place over a magnets for six min and the supernatant removed. The magnetic bead/antibody complexes were suspended in 2 ml ScintiVerse BD Cocktail (Thermo #163431). Vials were vortexed for 10 sec before quantitation by a Tri-Carb 2910 TR scintillation counter. An “[Inhibitor] vs. normalized response -- Variable slope” model (Graphpad Prism 7.0d) was fit to the METH serum standard curve data points to determine the IC_50_’s of each unknown serum sample.

### Disposition of METH after AAV-7F9-Fc Treatment

Mice given either 1 × 10^12^ vc/mouse 7F9-Fc therapy or vehicle were administered 3.1 mg/kg METH subcutanteously (sc). Thirty minutes later mice were sacrificed, and brains and trunk blood were harvested. After clotting, blood samples were centrifuged at 12,000 rcf for 10 min and serum was transferred into new tubes. Brains were homogenized in 4 ml of mass spectrometery grade H_2_O per gram of brain tissue.

A previously described extraction method for (3,4)-methylenedioxypyrovalerone was adapted for the extraction of METH and AMP in serum and brain tissue (Hambuchen *et al.*, 2017). Twenty-five microliters of each sample or standard was added to a 1.5 ml centrifuge tube. A volume of 125 μl ice-cold 30 ng/ml methamphetamine-D5 (internal standard) in acetonitrile was added prior to brief vortex mixing. After adding the internal standard solution to all samples and standards, the mixtures were vortex mixed for an additional 10 s prior to incubation for 10 min at 4°C. Afterward, the mixture was centrifuged at 20,000 rcf (at 4°C) for 5 min, and the supernatant was transferred to another 1.5 ml tube for drying in a Zymark TurboVap LV evaporator (SOTAX Corporation) under gentle nitrogen flow in a water bath at 40°C.

Dried samples and standards were reconstituted with 75 μl of 0.1% formic acid and vortex mixed for 20 s. After reconstitution, samples were centrifuged at 20,000 rcf (at 4°C) for 5 min, and 65 μl of supernatant was transferred to an autoinjector plate where 7.5 μl were injected onto an Acquity UPLC BEH C18 1.7 μm (2.1 i.d. × 100 mm) column (Waters Corp, Milford, MA) in an Acquity Ultra Performance Liquid Chromatography system connected to a Quattro Premier XE mass spectrometer (Waters Corp).

Analysis of unchanged METH concentrations in extracted samples were performed by a previously described liquid chromatography tandem mass spectrometry (LC/MS-MS) method (Hambuchen *et al.*, 2015). The lower and upper limit of quantification for METH and AMP was 1 and 1000 ng/ml, respectively. All predicted values for calibration and quality control standards were within ± 20% of the actual concentrations.

### Locomotor activity assays in mice

Locomotor studies were performed on the mice between days 21 and 37 post-AAV administration. The vehicle and treated mice (n=6/group) were administered either PBS (saline) or 1 × 10^12^ vc/mouse and allowed at least 21 days to express 7F9-Fc into the serum. Mice were allowed to equilibrate to their surroundings for 15 min before being administered PBS or 1.7, 3.1, 9.4 mg/kg METH sc. These doses were suppose to be prepared as 1.0, 3.0 and 10 mg/kg METH respectively, but afterwards were quantified by LC-MS/MS and found to be slightly different from the target doses. The corrected values were used for all data analysis. After drug administration, mice were placed back into their beam break boxes (Med Associates #MED-OFAS-MSU). Recording started once the mouse was placed in the box. Mouse activity was recorded for two hours post injection. Afterwards, mice were placed back into their home cages. There was at least a two-day washout period between testing sessions. Activity was expressed as total activity over 90 min post-administration of METH.

### Statistical Analyses

All data were analyzed using GraphPad Prism (version 7.0d, GraphPad Software, Inc.). Power analyses were performed to determine appropriate n values for each study using the http://clincalc.com/Stats/SampleSize.aspx site with power set to 0.8 and α = 0.05. Prior to analysis, any outliers were identified in the data using a Grubb’s outlier test with α = 0.05. Variances were assumed to be similar. The comparisons of the peak circulating concentrations of the construct between the different doses of AAV were tested with a one-way ANOVA using a Tukey’s correction for multiple comparisons when a significant difference was found between groups. Two-tailed, unpaired t-tests were used on the means of the IC_50_’s. Significance was determined for the locomotor assays with a two-way ANOVA using a Tukey’s correction for multiple comparisons when a significant difference was found between groups. The resulting data from the METH biodistribution study analyzed via 1-tailed, unpaired t-tests. Unless otherwise noted, significance was determined at p < 0.05 and denoted with an astrisk.

## Results

### Determining Optimal AAV Dose for 7F9-Fc serum expression

A dose-dependent relationship was observed in Figure 2A between the dose of AAV-7F9-Fc administered and the circulating concentration of anti-METH antibodies. Over the observed 170 day period, the mice administered 1 × 10^10^ vc/mouse peaked at 48 μg/ml, those administered 1 × 10^11^ vc/mouse peaked at 1,785 μg/ml, and those administered 1 × 10^12^ vc/mouse peaked at 3,831 μg/ml. A 1-way ANOVA of the peak values yielded significant variation between groups F(2, 12) = 2.058, p<0.0001. Tukey post-hoc tests showed there were significant differences in expression between each of the observed peak circulating concentrations (p<0.05).

**Figure 2:**
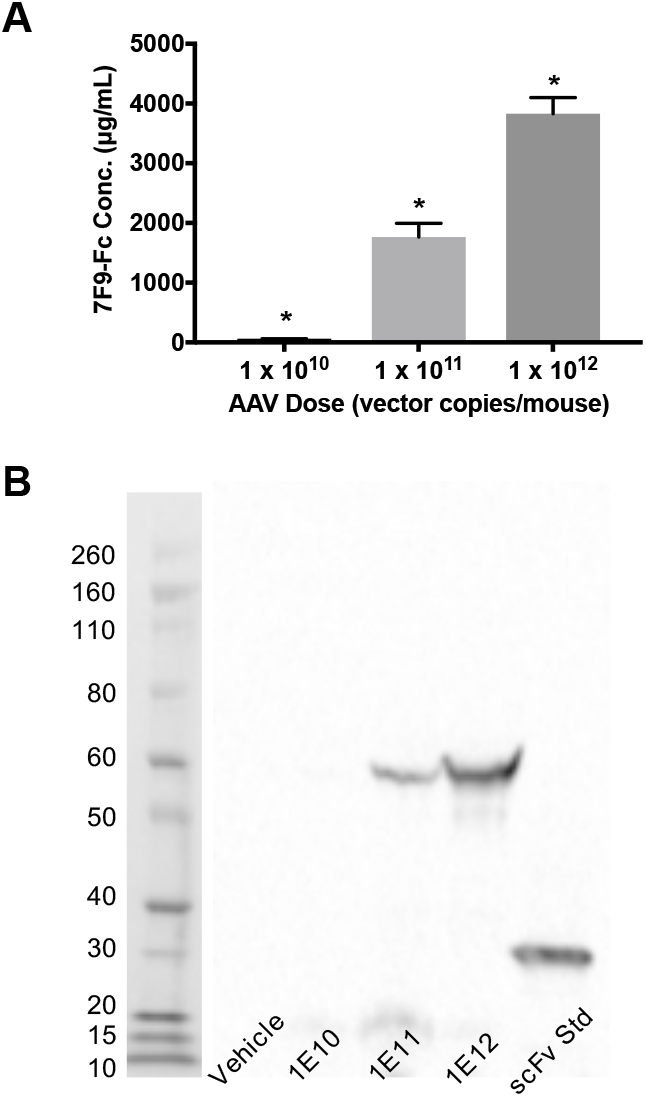
A dose-dependent relationship exists between the dose of AAV given to mice and the level of expression of 7F9-Fc. A & B: Functional ELISAs were performed on mice serum. A: Peak serum concentration of 7F9-Fc. (*, p<0.05) B: A western blot comparing the expressed size of construct (61 kDa) to that of a predecessor scFv standard of know size (23 kDa).

### Biocharacterization

Serum samples taken from mice on day 22 post-AAV administration were pooled by AAV dose (vehicle, 1 × 10^10^, 1 × 10^11^, or 1 × 10^12^ vc/mouse). The intact monomeric 7F9-Fc bands appeared near the 60 kDa molecular marker band, as expected. The 7F9-Fc protein monomers were calculated to be 61.6 kDa. As observed in Figure 2B, the 7F9-Fc bands appear to increase in intensity, but the band for the lowest concentration is not observable. This is supported by the very low 7F9-Fc concentrations of the 1 × 10^10^ dose reported in Figure 2A.

### Affinity

Pooled serum samples from mice expressing 7F9-Fc collected 22 days after AAV administration showed the IC_50_ value for 7F9-Fc was 17 nM, while the mAb7F9 IC_50_ value was of 32 nM (Figure 3) (Stevens, Tawney, *et al.*, 2014).

**Figure 3:**
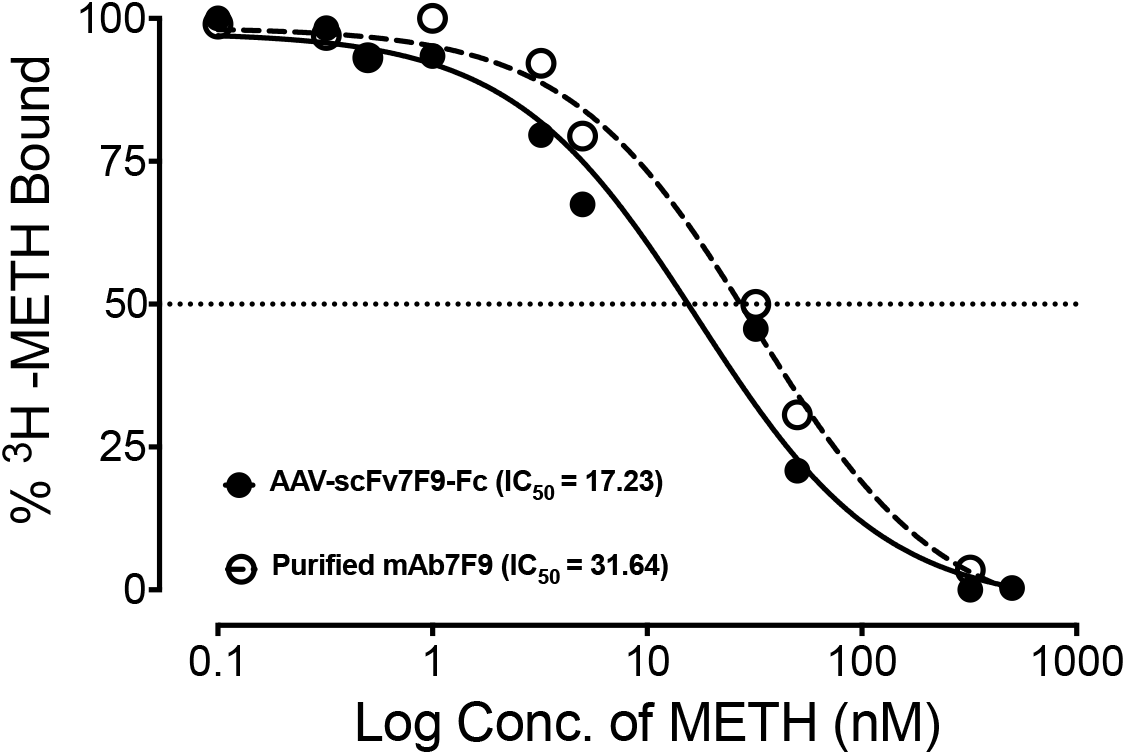
Comparison of IC_50_ values for METH between culture-produced mAb7F9 and mouse-expressed 7F9-Fc. The IC_50_ (nM) for all variants was estimated at 50% ^3^H-METH bound (dotted line). Data shown as mean ± SEM, n = 8 per group.

### Duration 7F9-Fc Expression in Mice

After selecting the 1 × 10^12^ vc/mouse dose of AAV-7F9-Fc for futher studies, a second set of mice were administered the dose via tail vein to determine the duration of 7F9-Fc expression and the expression was followed over an approximately 8 month period.. As can be observed in Figure 4, the mice expressed 7F9-Fc anti-METH antibodies for at least 239 days after a single AAV administration. During this nearly eight-month period, the mice exhibited an average circulating concentration of 2,116 ± 1,543 μg/ml, with a peak of 5,228 ± 1,637 μg/ml on day 63 post-AAV administration.

**Figure 4:**
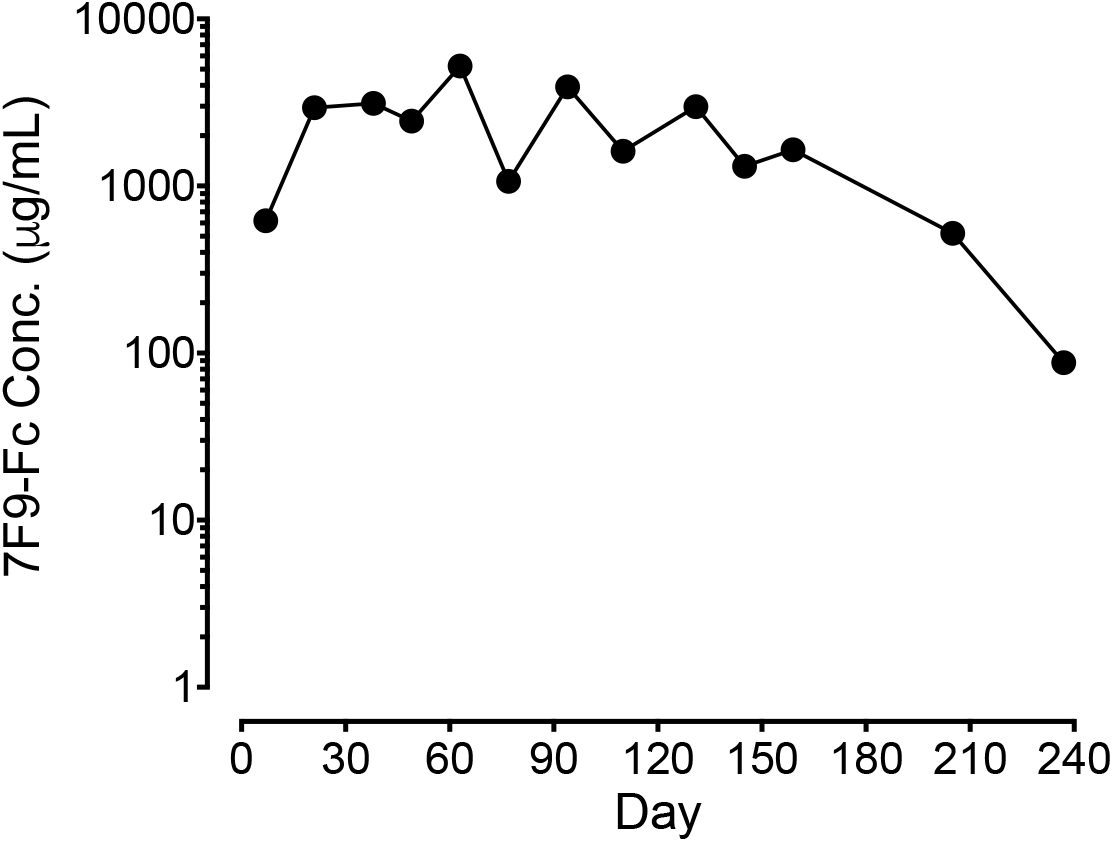
7F9-Fc expression persists over 8 months. Mice given 1 × 10^12^ vector copies/mouse showed expression for at least 8 months, with an average expression of 2,100 ug/ml. Functional ELISAs were performed on mouse serum. Data given as mean ± SEM, n = 6 per group.

### Locomotor Studies

In all experimental groups, administration of saline had no systematic effects on mouse behavior, and locomotor activity remained low for all subjects. In the vehicle group, administration of METH increased the total distance traveled during a 90 min period in a dose-dependant manner, with mice treated with 3.1 and 9.4 mg/kg METH exhibiting significantly greater locomotor stimulation than when injected with saline [F(4,16)=8.474, p=0.0007]. No stereotypy was observed at any concentration of METH in the vehicle mice. The mice given the AAV-7F9-Fc therapy did not exhibit a significant increase in basel horizontal distance traveled regardless of the METH dose administered [F(4,16)=8.474, p=0.0007]. No stereotypy was observed at any concentration of METH in the treated mice. A tukey post-hoc test showed significantly lower total movement in a 90-min period after being administered of either 3.1 or 9.4 mg/kg METH sc in mice given the AAV-7F9-Fc therapy than the mice given vehicle as shown in Figure 5 (p<0.05 for both).

**Figure 5:**
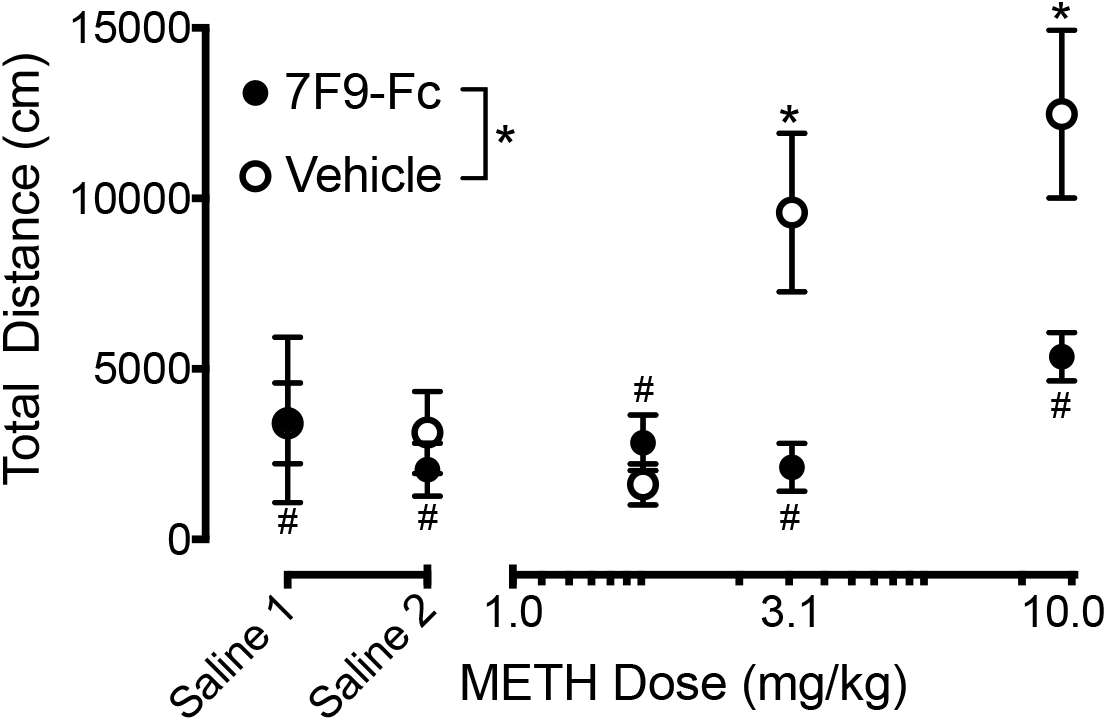
7F9-Fc significantly reduces the psychostimulant effects of METH at 3.1 and 9.4 mg/kg METH doses. Two groups of mice, vehicle and mice given 1 × 10^12^ vc AAV-7F9-Fc/mouse, were either administed saline or increasing doses of METH sc. The locomotor activity measured as total distance traveled was recorded for 90 min. Comparison between AAV-7F9-Fc treated mice (#, p < 0.05). Comparison between vehicle and AAV-7F9-Fc mice (*, p < 0.05) Data are shown as mean ± SEM, n= 6 per group.

### Physiological Efficacy

Four months post-administration of either AAV or vehicle, the same mice used in the locomotor assays were given 3.1 mg/kg METH sc. As can be observed in Figure 6A, mice treated with AAV-7F9-Fc showed 1.2 times lower METH in their brains (2407 ng METH/g brain) than vehicle mice (2863 ng METH/g brain). Further, the treated mice showed over 2.6 times more METH in their serum (828 ng METH/ml serum) than vehicle mice (324 ng METH/ml serum) (Figure 6B).

**Figure 6:**
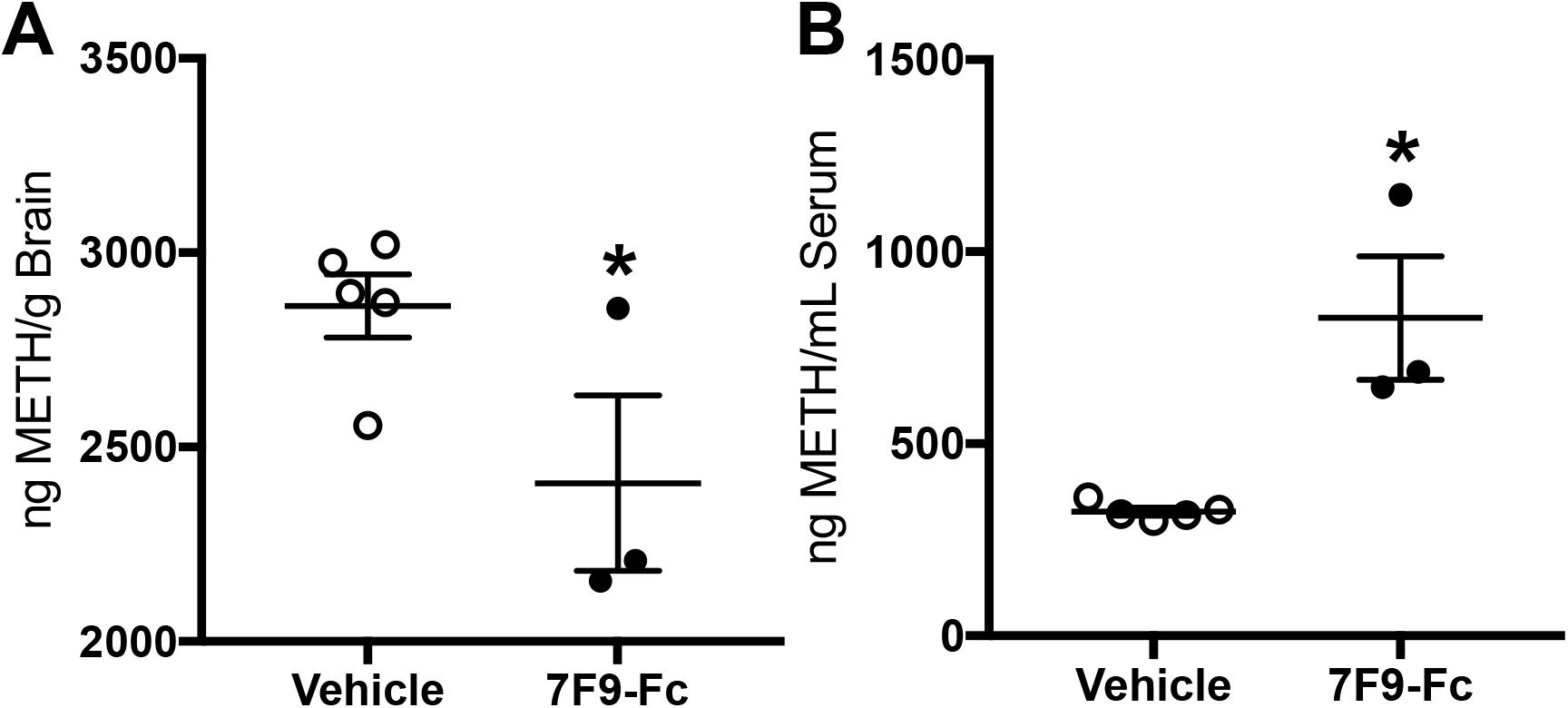
A comparison of METH brain and serum concentrations after a 3.1 mg/kg *sc* injection of METH. Two groups of mice, vehicle and mice given 1 × 10^12^ vc AAV-7F9-Fc/mouse, were administered 3.1 mg/kg METH sc. Thirty minutes after METH administration, mice were sacrificed and brains (A) and serum (B) were collected. METH concentrations were determine by LC/MS-MS. Data points are shown as mean ± SEM, n = 3-5 per group. An astrisk indicates the values are significant at p < 0.05.

In both this study (data not shown) and as previously reported with the same concentration of AAV, no lasting adverse effects were observed as a result of the high AAV dose (Hay *et al.*, 2018).

## Discussion

The purpose of this study was to design and test in vivo and ex vivo, a long-lasting AAV-delivered anti-METH antibody therapy. The overarching hypothesis was that continuous expression of a high affinity anti-METH antibody with a persistent serum residence time could be a viable approach to creating medications to aid in treatment of METH use disorders. We have previously presented proof-of-concept data on expression of such an approach with an anti-METH mAb transgene (Hay *et al.*, 2018). Here, we wanted to increase the transgene expression and provide data on its functional capacity against METH effects. For this purpose, we engineered a single gene sequence based on a high affinity monoclonal antibody (mAb7F9) that could efficiently be packaged into AAV8 particles. By incorporating an Fc region into the construct, detectable expression lasted at least another month with average expression levels of 2,100 μg/ml – thirty times greater than the first generation constructs, (data not shown), likely due to significantly reduced clearance of scFv-7F9-Fc (Hay *et al.*, 2018).

For this study, we used three different concentrations of the AAV-7F9-Fc therapy: 1 × 10^10^, 1 × 10^11^, and 1 × 10^12^ vc/mouse to determine dose-response relationships between the dose of AAV-7F9-Fc and concentrations of expressed scFv-7F9-Fc. In addition to providing evidence of proper secretion of the 7F9-Fc protein, Figure 2B also shows increasing concentrations of the 7F9-Fc protein in mouse serum collected 22 days post-AAV administration. Because a significantly larger circulating concentration of our anti-METH construct at the 1 × 10^12^ vc/mouse dose was detected, we chose to use the 1 × 10^12^ vc/mouse dose to determine the the potential upper limit of reduction of the pharmacological effects of METH. Future studies will be required to determine what dose of the AAV-7F9-Fc therapy provides the optimal circulating concentrations for the AAV dose given.

The results from the locomotor study suggest that our anti-METH therapy is able to mitigate the psychostimulant effects of METH at relatively high doses. As can be observed in Figure 5, the 9.4 mg/kg METH dose for the treated mice, while not significantly greater than the saline doses, appears to be visually greater than the two lower METH doses. Because this visible increase in total distance traveled is roughly shifted to the right by about one log value, this could suggest that the AAV-7F9-Fc therapy decreases METH’s potency by about one log value. While a more extensive panel of behavioral assays would be required to confirm, the data appears to suggest that the AAV-7F9-Fc treated mice are beginning to exhibit psychostimulant effects of METH around the 9.4 mg/kg METH dose, compared to the vehicle mice beginning to exhibit the psychostimulant effects of METH between the 1.7 and 3.1 mg/kg METH doses. The 1.7 and 3.1 mg/kg METH sc doses were chosen because our lab group has used these doses to model moderate to high clinically relavent human doses (Gentry *et al.*, 2004, 2009). Because the circulating concentrations of 7F9-Fc were observed to be so high, the 10 mg/kg METH sc dose was chosen to model extremely high doses of METH (Cruickshank and Dyer, 2009; Courtney and Ray, 2014). The reduction of the psychostimulant effects observed in these locomotor studies should persist throughout the expression period of 7-8 months in mice, and possibly longer in humans.

As previously noted, other groups (Hicks *et al.*, 2012; Rosenberg *et al.*, 2012, 2013) have successfully expressed full IgG antibodies using gene transfer. When designing our new construct, we observed that one accepted approach was to connect both the heavy and light chains of the IgG together with a furin and 2A cleavage sequence (Fang *et al.*, 2005; Ho *et al.*, 2013). The furin sequence once translated, could recognize and cleave apart the 2A sequence and the light and heavy chains of the antibody could assemble and dimerize. While this approach is viable, we hypothesized that a simpler approach might result in higher circulating concentratins due to less cellular processing. The scFv-Fc fusion construct can be expressed as a single continuous chain, which does not require post-translational cleavage steps for assembly (Unverdorben *et al.*, 2016). Further, scFv-Fc fusion constructs were previously shown to have similar half-lives compared to full IgG antibodies, primarily due to similar sizes and utilizing the FcRn pathway, which greatly increases the half-life of proteins (e.g., albumins and antibodies) beyond what would be predicted for proteins of their size (Ghetie *et al.*, 1996).

We also chose to utilize the IgG_2_ subclass rather than the IgG_1_ subclass as others have (Hicks *et al.*, 2012; Rosenberg *et al.*, 2012). The IgG_2_ subclass has a similar affinity to the FcRn, but IgG_2_ has a lower affinity for the Fc receptors on effector cells and is far less effective at activating complement, which is not needed in our treatment modality (Vidarsson *et al.*, 2014). These differences could translate to a decrease in undesirable systemic immune responses as a result of the expressed anti-METH antibodies.

The affinity of 7F9-Fc to METH was found to be 17 nM and the affinity of the purified standard mAb7F9 was 31 nM. Although, the variable region for both of these proteins is the same, the remaining portions of the protein are different, which could explain the minor difference in affinity *17 nm vs 32 nm). This pattern was observed by the scFv constructs of scFv7F9 showing an affinity to METH of 22 nM compared to an affinity of 48 nM of the purified standard (Hay *et al.*, 2018). This is the first time our group has compared the affinity of the 7F9-Fc construct to the original IgG mAb7F9.

The biodistribution study (Figure 6) suggests that the AAV-7F9-Fc therapy is able to significantly lower the amount of METH in the brain and sequester the METH in the serum. After administering a 3.1 mg/kg METH sc dose, and collecting brain and serum from the mice 30 min post-METH administration we observed 1.2 times lower METH concentrations in the brains of treated mice and an over 2.5 times greater concentration of METH in the serum of the treated mice compared to the vehicle mice. This indicates that 30 min post-METH administration, approximately T_max_ for METH in mice brain and serum (Magyar *et al.*, 2007), significantly greater concentrations of METH are being sequestered in the serum of the treated mice. This study was performed on day 237 (last point in Figure 4), the antibody circulating concentration of 87 μg/ml resulted in a METH concentration of 827±279 ng/ml. This resulted in a METH:7F9-Fc molar ratio of 1,561:1 (5.54 mM METH and 3.55 μM 7F9-Fc), but even with this low antibody concentration, we observed sequestration of METH in the serum and reduction in brains of the treated mice compared to the vehicle mice. Had the assay been performed shortly after the mice had finished the locomotor assay with concentrations of about 600 μg/ml (data not shown), the reduction of METH in the brains would likely have been much greater. This dose of METH was selected to challenge the circulating antibodies and it shows that even after the antibody is surmounted, physiological efficacy can be observed. In comparison, one minute after a 0.8 μg nicotine dose, Hicks *et al.* (2012) observed 7.2 times greater amounts of nicotine in the serum, and three minutes after a 0.25 μg cocaine dose, Rosenburg *et al.* (2012) observed a 10-fold increase in serum cocaine levels compared to vehicle mice.

In conclusion, these studies show the design, expression, and pre-clinical characterization of a high affinity, long duration of action anti-METH antibody-based gene therapy. The incorporation of an Fc region resulted in sustained circulating concentrations of over 2.1 mg/kg in mice over a period of eight months. Initial biodistribution characterization suggests that the therapy was capable of sequestering METH in the serum and out of the brain, even at lower circulating concentrations. Data from behavioral assays suggest that this anti-METH gene therapy is able to prevent mice from exhibiting METH-induced psychostimulant effects after relatively large subcutaneous METH doses. Taken together, these intial studies suggest that a scFv-Fc based AAV therapy can prevent the psychostimulant effects of METH by reducing the amount of METH able to enter the brain in mice.

## Acknowledgements

We would like to thank Melinda Gunnell, and Robin Mulkey for their expert technical assistance.

## Authorship Contributions

Participated in research design: Hay, Peterson

Conducted experiments: Hay, Ewing, Hambuchen, Zintner, Small, Bolden

Contributed new reagents or analytic tools: Peterson, Owens, Fantegrossi, Margaritis

Performed data analysis: Hay, Hambuchen, Peterson

Wrote or contributed to the writing of the manuscript: Hay, Ewing, Hambuchen, Zintner, Small, Fantegrossi, Margaritis, Owens, Peterson

## Footnotes

The project described was supported by the National in Insitutes of Health (NIH): National Institute on Drug Abuse (NIDA): R01-DA036600, the National Institute for General Medical Sciences:T32-GM106999 and R25-GM083297.

